# Air-nanobubbles ineffective to reduce pathogenic bacteria in fresh and brackish waters

**DOI:** 10.1101/2021.08.27.457885

**Authors:** Jose A. Domingos, Qianjun Huang, Hong Liu, Ha Thanh Dong, Nareerat Khongcharoen, Phan Thi Van, Nguyen Huu Nghia, Pham Thai Giang, Pham The Viet, Sophie St-Hilaire

**Affiliations:** City University of Hong Kong, 5/F, Block 1, To Yuen Building, 31 To Yuen Street, Kowloon, Hong Kong; Food Inspection and Quarantine Technical Centre, Futian, Shenzhen, Guangdong, China; Faculty of Science and Technology, Suan Sunandha Rajabhat University, 1 U Thong Nok Rd., Dusit, Bangkok 10300, Thailand; Research Institute for Aquaculture No.1, Dinh Bang, Tu Son, Bac Ninh, Vietnam

**Keywords:** *Aeromonas* spp., nanobubbles, *Vibrio* spp., water treatment

## Abstract

Nanobubble (NB) technology has been hailed as a novel way to disinfect water. Previous studies suggested that when NBs collapse, they create shock waves that result in OH^-^ free radicals, which can damage cells, including bacteria. In this study, we investigated, through a series of 11 experiments, the potential use of air nanobubbles (128 ± 44 nm, mean ± SD) to reduce the concentration of various pathogenic bacteria including *Aeromonas hydrophila, A. veronii, Vibrio parahaemolyticus*, and *Streptococcus agalactiae* under controlled, tank-based laboratory conditions. Despite the high number of nanobubbles continuously added to a relatively small volume of water in experimental tanks (50-100 L), we did not observe a consistent or significant decrease in bacteria that would control disease outbreaks. Although most of the experiments were conducted in fresh water on *A. hydrophila*, results were consistent across fresh and brackish water experiments, Gram-negative and Gram-positive bacteria, and a range of nanobubble concentrations. This study suggests air nanobubbles on their own are inadequate to significantly reduce high levels of pathogenic bacteria in water. We propose to explore other gases for improving the disinfection properties of this technology.

**SIGNIFICANCE STATEMENT:** Air nanobubbles did not sufficiently reduce the level of bacteria in laboratory experiments.

## 1. INTRODUCTION

Nanobubbles (bubbles <1µm in diameter) have several characteristics, which are hypothesized to confer unique disinfection properties (Agarwal et al., 2011). Their long residence time in water is attributed to their low buoyancy (Agarwal et al., 2011; Gurung et al., 2016) and their high zeta potential prevents bubble aggregation (Michailidi et al., 2019). The high internal pressure within the nanobubbles permits efficient gas transfer, and their large surface area to volume ratio promotes chemical reactions (Gurung et al., 2016). As nanobubbles lose gas to the surrounding water the internal pressure in the bubble increases, eventually when nanobubbles collapse they generates small shock waves, which create hydroxyl radicals (HO·) (Agarwal et al., 2011). Free radicals oxidize and damage cell membranes removing electrons (Halliwell, 1996). Varying the gas compositions of the nanobubbles can increase their stability, oxidizing potential, and electric polarization, which further improves their disinfection capability (Gurung et al., 2016; Tatek et al., 2017).

There are only a few published studies on the use of nanobubble technology in aquaculture and the majority of these examine the disinfection properties of ozone nanobubbles (Kurita et al., 2017; Imaizumi et al., 2018, Jhunkeaw et al., 2020). For aquaculture in particular, ozone may be used to disinfect and clarify water; however, its use has a number of practical limitations. Ozone is highly toxic to all life forms, including the animals being farmed and the workers if inhaled. It is also highly corrosive to the equipment it comes in contact with. In water containing bromide (Br^-^), such as sea water, the use of ozone generates other undesirable toxic compounds (i.e. ozone-produced oxidants) such as bromine (Br_2_) and bromate (BrO_2_^-^) (Tynan et al., 1993, Schroeder et al., 2014); therefore, ozone disinfected water needs to be monitored and toxic compounds need to be removed before it can be delivered to tanks with live animals.

Furthermore, ozone is expensive to produce in large quantities, as its generation requires controlled temperature and humidity conditions and, in a practical aquaculture setting with variable levels of organic matter, it is difficult to dose accurately (Mundya et al., 2018).

As nanobubble technology is still in its infancy, the potential bacteriostatic or bactericide properties of nanobubbles generated with innocuous gases, such as air, is yet to be determined. Therefore, in this study we report on several experiments, conducted by multiple research groups, to investigate the disinfecting potential of air nanobubbles against pathogenic bacteria of concern to cultured fish and shrimp in fresh and brackish waters under controlled laboratory settings.

## 2. MATERIAL AND METHODS

### 2.1 Characterization of nanobubbles

We generated nanobubbles using a commercial nanobubble generator aQua+075MO (AquaPro Solutions Pte Ltd, Singapore). To verify the production of nanobubbles we used a nanoparticle analyser NanoSight LM10 (Malvern Panalytical Ltd), which measured particle size and concentration in the water. As the NanoSight LM10 measures particles and it cannot distinguish between gas bubbles and solid particles, e.g. dust we produced bubbles in distilled water to minimize nanoparticles and estimate the quantity of nanobubbles produced by the nanobubble machine. To reduce the noise from nanoparticles we cleaned and rinsed the equipment with distilled water prior to conducting the study. Control samples were taken at time zero, immediately before the nanobubble generator was switched on, to account for any particles within the system. The nanobubbler was run in 30 L (Trial 1, replicate 1) and 12 L (Trial 2, replicates 2 and 3) of distilled water for 20 min, after which several 40 ml water samples were collected in 50 ml sterile Falcon tubes and kept at room temperature with closed lids until analysis up to three weeks later. To assess bubble decay characteristics over time, samples were analysed at distinct time intervals, up to three weeks after collection. To avoid contamination, each water sample was tested only once, i.e. after the tube was opened and triplicate sample readings were taken, the tube was discarded.

### 2.2 Effects of air nanobubbles on bacteria

To investigate the lethal potential of air nanobubbles on pathogenic bacteria in fresh and brackish water, 12 experiments were conducted across three countries by three separate teams (Table 1). We report our data together in this manuscript to increase the external validity of our non-significant findings vis-à-vis the effects of air nanobubbles on high concentrations of bacteria. Experiments 1 through 7 were carried out at the Food Inspection & Quarantine Technical Center (FICS) of Shenzhen, China; experiments 8, 9a, 9b, and 11 were carried out at the Suan Sunandha Rajabhat University in Bangkok, Thailand, and experiment 10 was carried out at the Research Institute of Aquaculture (RIA1) in Hanoi, Vietnam. Four bacterial species known to cause mortalities and severe economic losses in cultured fish and shrimp, namely *Aeromonas hydrophila, A. veronii, Vibrio parahaemolyticus* (Gram-negatives), and *Streptococcus agalactiae* (Gram-positive) were either exposed to air nanobubbles (treatments) or not. Our controls were provided gentle aeration using an air stone for the same length of time as the longest nanobubble treatment. In one experiment (Experiment #3), the bacterial pathogen used was not identified accurately, but the data were included for completeness.

**TABLE 1.**
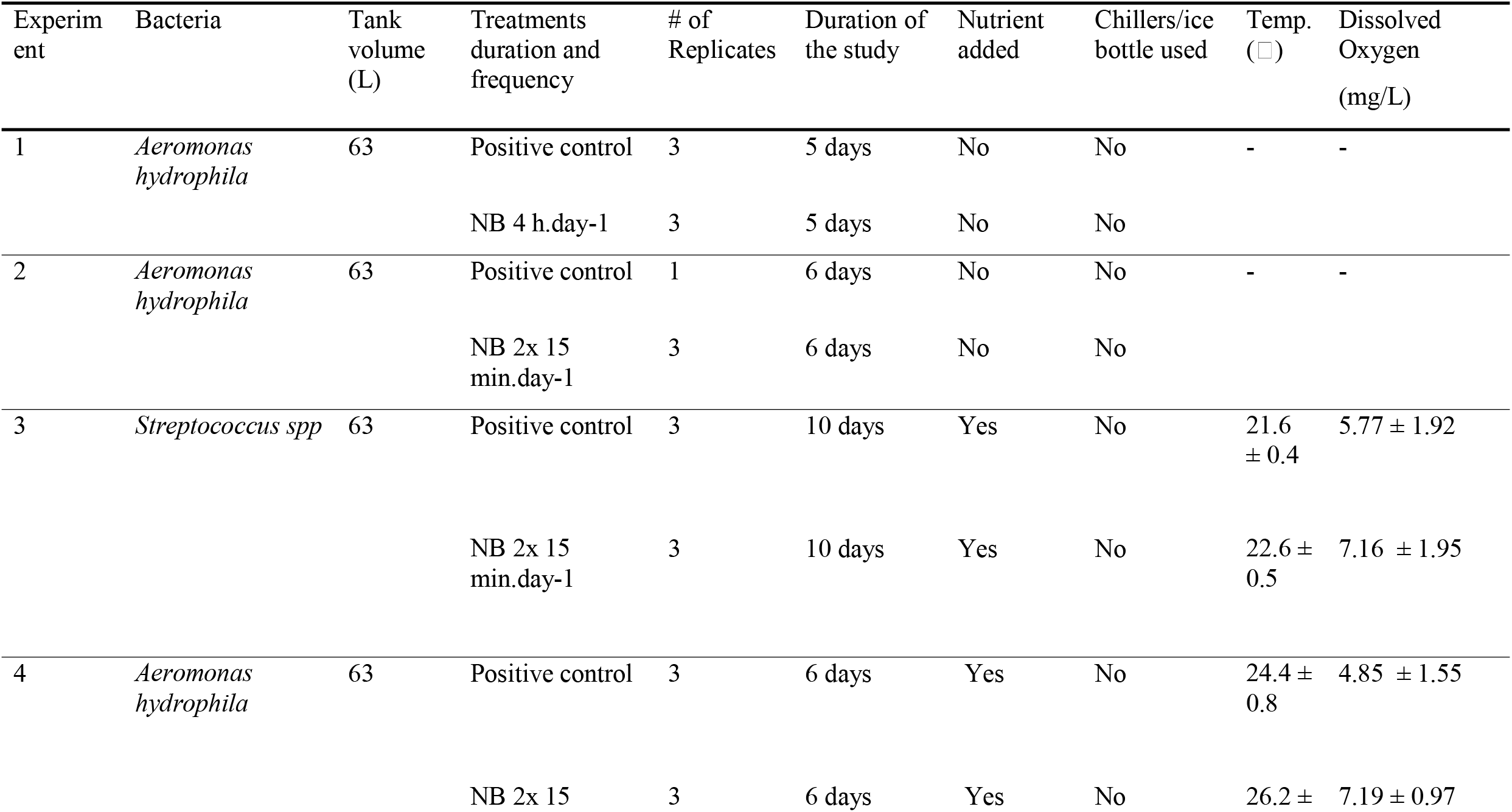
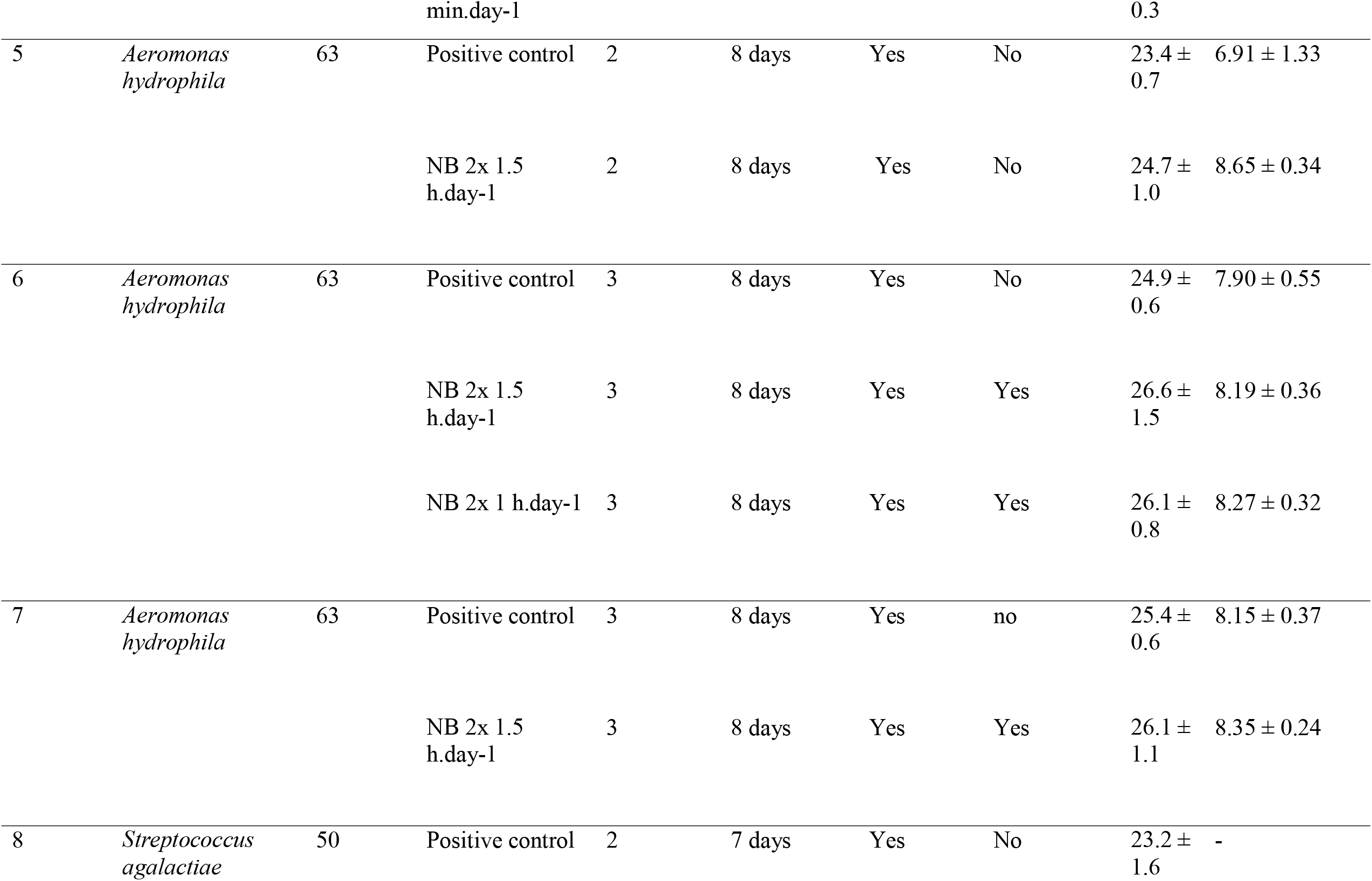
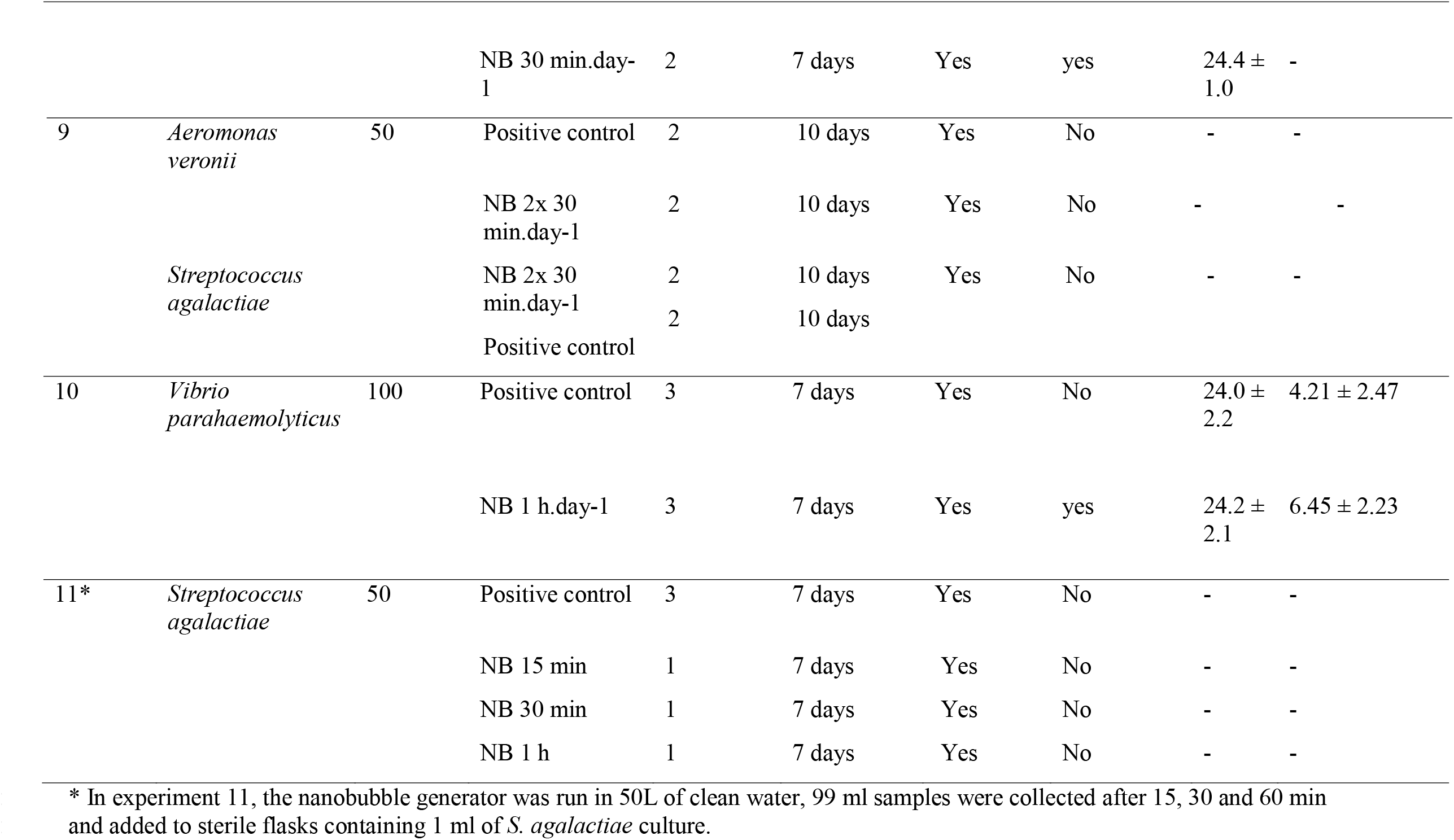
Description of experiments to assess the potential disinfecting effects of air and oxygen nanobubbles (NB) against diverse bacteria pathogenic to fish and shrimp previously inoculated in water of experimental tanks (except for experiment 11). The experiment with *Vibrio parahaemolyticus* (10) was conducted in brackish water (15 ppt); all others were conducted in freshwater. Temperature and dissolved oxygen values represent mean ± standard deviation.

Bacterial strains were sourced from each institution’s collection, and cultures were grown in sterilized tryptic soy broth (TSB) or Nutrient Broth (NB), for *V. parahaemolyticus*. With the exception of experiment 11, which was a flask experiment, experimental tanks were inoculated with a single dose of bacteria aimed at a concentration of 10 ^6^ to 10 ^7^ cfu·ml-1 on “day 0” (pre-treatment). Tanks in different laboratories were between 50 and 100 L in size. De-chlorinated tap water was used for all experiments. In experiments with *Vibrio parahaemolyticus*, brackish water was made by dissolving sea salt in de-chlorinated tap water at a concentration of 15 parts per thousand (ppt). All nanobubble exposure ranged between 15 min to 3 hr per day over 3 to 10 consecutive days, depending on the experiment (Table 1). Nanobubble treatments in experimental tanks were initially restricted to 15 min periods in order to avoid overheating the water, because the nanobubble generator increased water temperature at a rate of approximately 0.75 □ per 10 min in 75 L tanks under experimental conditions (data not shown).

In order to assess longer nanobubble treatment duration, experimental tanks were individually connected to chillers. In centers where chillers were not available (4 separate trails) frozen ice water bottles were added to tanks in order to maintain a stable water temperature, comparable to the control tanks (Table 1). Further, to avoid the natural decrease in bacterial counts seen during early trials, 40 or 50 ml of TSB was added to the tanks in some of the later experiments (Table 1). The addition of TSB was usually done on alternating days starting on day 3. In the case of Experiment 10 the researchers added TSB every day starting on day 2.

For experiment 11, the first study we did on the Gram-positive bacteria *S. agalactiae*, we exposed the bacteria to a single dose of nanobubbles without running the bacteria through the nanobubble generator. This experiment was conducted in 100 ml flasks, which were filled with 99ml of nanobubble treated de-chlorinated tap water and 1 ml of bacterial culture. Non-treated, de-chlorinated water was used instead of the nanobubble-exposed water for the control group. Three different concentrations of nanobubbles were used for this experiment to create a dose effect: 50 L of stock water was treated through the nanobubble generator for 15, 30, and 60 min to generate the 99 ml of nanobubble treated water. We only used one flask for each treatment group.

To monitor treatment effect, water samples were collected from each tank, or flask in the case of experiment 11, every day or every other day. Serial dilutions (1:10) were made and water was plated in duplicate on different media, depending on the pathogen. *A. hydrophila* was plated in brain heart infusion agar (BHIA), *V. parahaemolyticus* was plated nutrient agar with 2 % NaCl, and *S. agalactiae* and *A. veronii* were plated in tryptic soy agar (TSA). Bacteriology results were report as colony forming units (cfu) ·ml-1, which was determined from the serial dilution plates where there were discrete quantifiable numbers of colonies on the plates (i.e. between 30 to 200 colonies per plate). We provide raw data on cfu·ml-1 for each experiment graphically to illustrate trends in bacterial concentration across time in different experiments.

To standardize the bacterial levels across tanks and experiments we calculated the change in cfu.ml-1 between two day intervals. Because not all experiments measured the bacterial concentration every day, we assessed the change in the bacterial concentration every two days by the following formula:

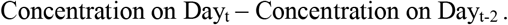

Eight out of 10 laboratory tank experiments had a crash in bacteria in both the control and the nanobubble treated tanks on or shortly after day 4 so we only analysed whether there was a significant difference in the change in bacterial concentration at the start of the study (i.e. first two days). To assess whether the difference in the relative change in bacterial concentrations between day 0 and day 2 was statistically different controlling for the experimental effect we conducted a nested general linear model in Minitab 19 (Minitab, Inc., Statistical Software (2010), State College, PA). The treatment was nested within each experiment (n=11). We had 11 experiments in our statistical analysis because we divided experiment 9 into two studies based on the bacteria used. Different exposure times to nanobubbles (30 min to 3 hr) and different pathogens were not assessed separately for this analysis due to the limited data. We assessed the standardized residuals for equal variance and the distribution of the standardized residuals was visually inspected for normality. Because our sample size was relatively small and we could not meet the parametric assumption of normally distributed residuals, we also evaluated the difference in the bacterial concentration between day 2 and day 0 for controls and treated tanks for each experiment separately using the non-parametric Kruskal-Wallis test.

In addition, water temperature and dissolved oxygen were monitored daily with a digital probe (YSI, Yellow Springs, OH, USA). Overall average temperature and range of temperatures in each experiment are summarized in Table 1.

## 3. RESULTS

### 3.1 Characterization of nanobubbles

Data from the Nanoparticle analyser readings revealed that nanobubbles were stable both in size (diameter of 128 – 154 nm) and concentration, over a three-week period (Figure 1A). The concentration of bubbles in distilled water only declined from 88 to 54 million·ml-1 during the three-week period (Figure 1B). Data from the second trial confirmed the bubble size produced by the nanobubble generator had a mean diameter of 128 ± 44 nm (SD) (Figure 1C). No significant changes in the size of the bubbles were observed after one week (Figure 1D). Although a small quantity of particles was detected in non-treated (control samples), the concentration represented only 0.8 % of the particles detected after exposure to the nanobubble generator. After 20 min of operation, an average of 9 × 10^7^ and 1.2 ×10^8^ nanobubbles.ml-1 were produced in 30L (Trial 1) and 12 L (Trial 2) of water, respectively. In both of these trials, 50 % of the particles (bubbles) measured were equal to or smaller than 112 nm and 69 % were below 150 nm in diameter.

**FIGURE 1.**
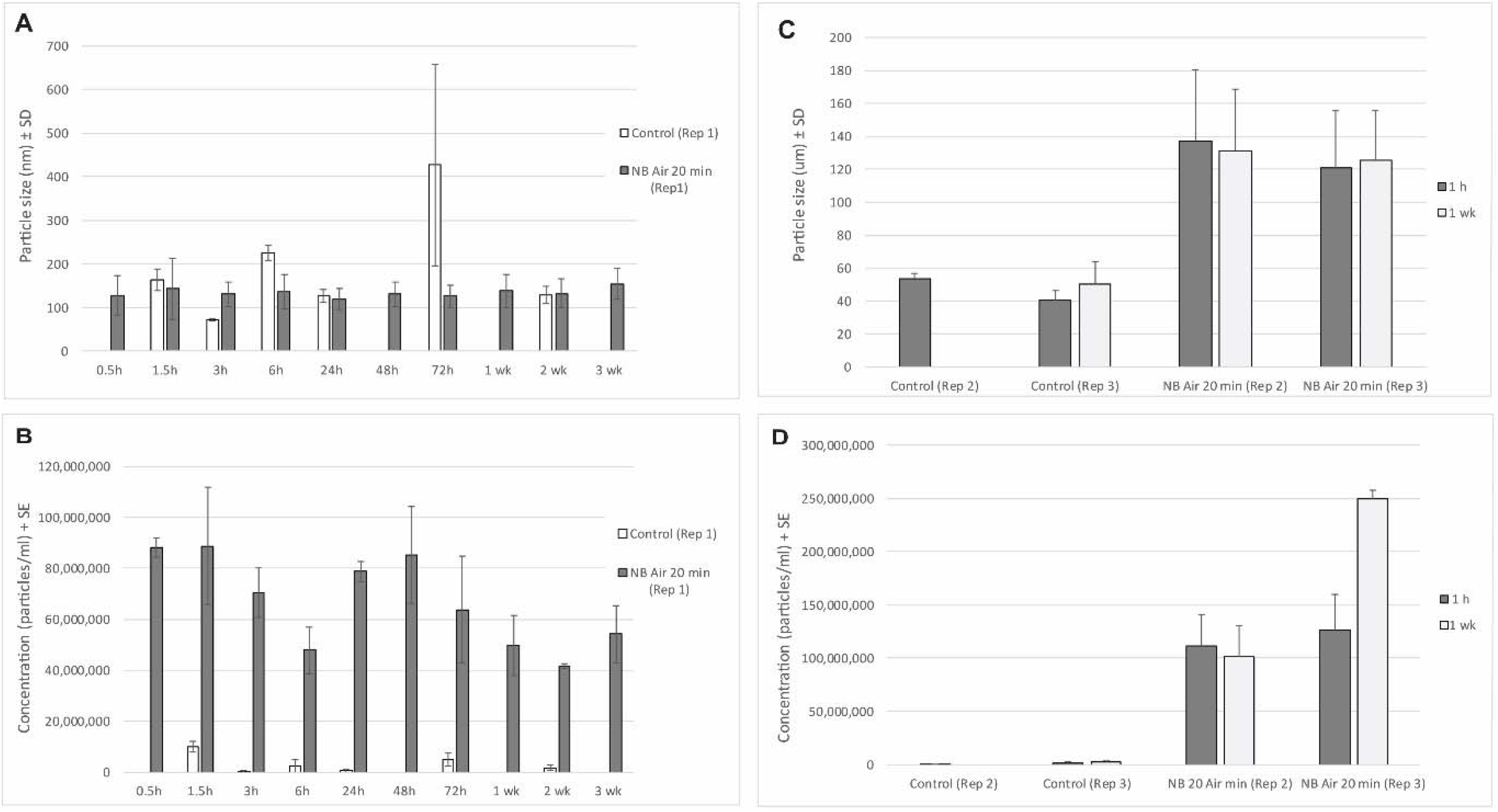
Particle size (A, C) and concentration (B, D) in distilled water before (Control, background particles) and after the nanobubble generator was run for 20 min (NB Air 20 min).

### 3.2 Effects of air nanobubbles on bacteria

We observed varied effects of air nanobubbles on bacterial colony forming units per ml (cfu·ml-1) (Figure 2). Seven out of 11 experiments (experiment # 3,4,7,8,9a,9b, and 10) had an increase in bacteria concentration over the first few days of the application of nanobubbles in both the NB exposed tanks and the control tanks (Figure 3). Four experiments (Experiment # 1, 5, 6, and 7) had a decrease in bacteria cfu·ml-1 in the tanks exposed to NB within the first few days of the experiment (Figure 3), but this was either also observed in the control tanks (Exp. #1) or was only a relatively small (non-significant) difference in the bacterial concentration (i.e. 10 to 20-fold reduction between control and experimental tanks (Figure 2).

**FIGURE 2.**
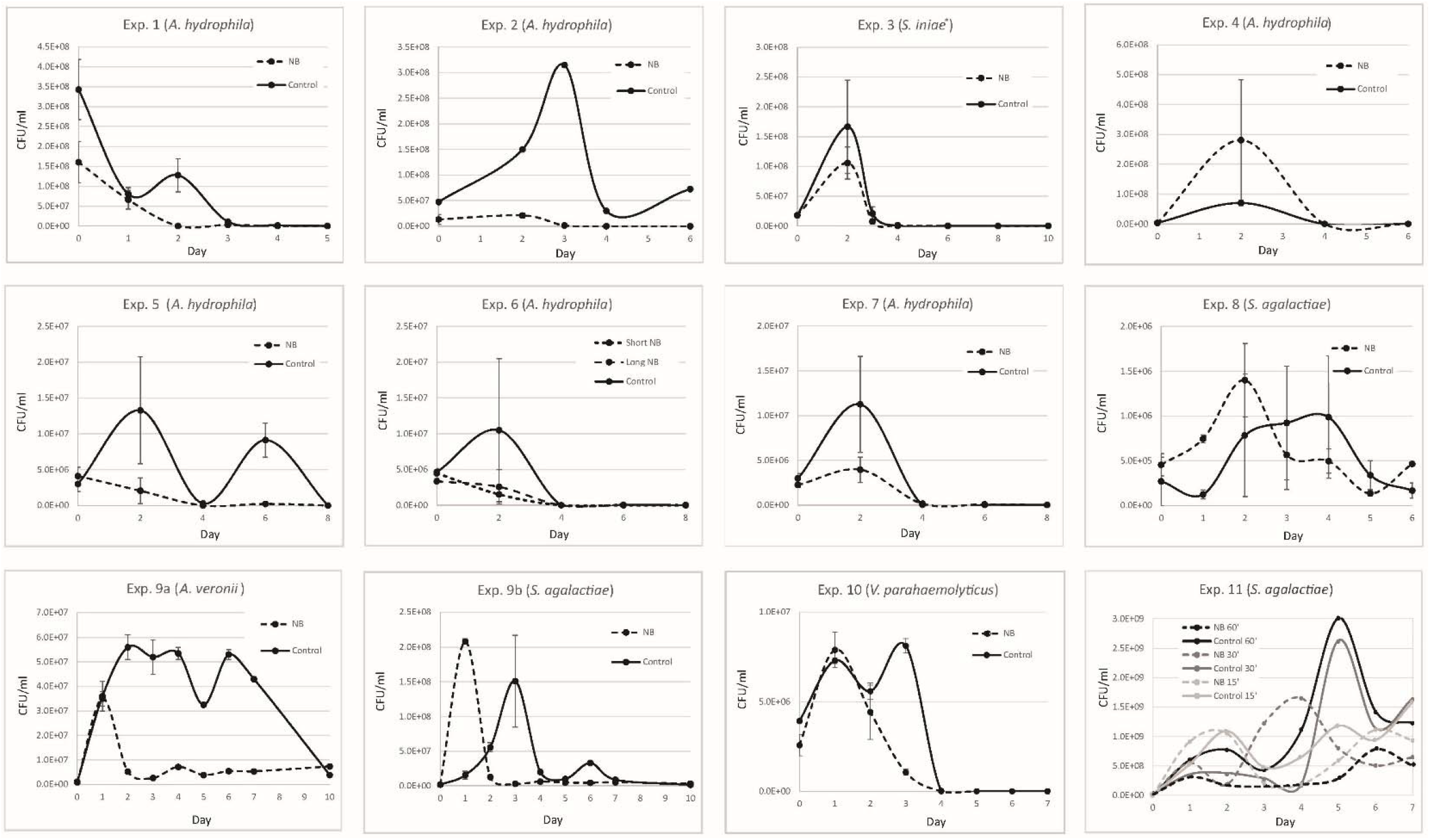
Concentration of bacteria in cfu.ml-1 over time for all experiments included in the study. Experiment 11 was conducted in flasks. Experiment 2 did not have replication but is included for completeness.

**FIGURE 3.**
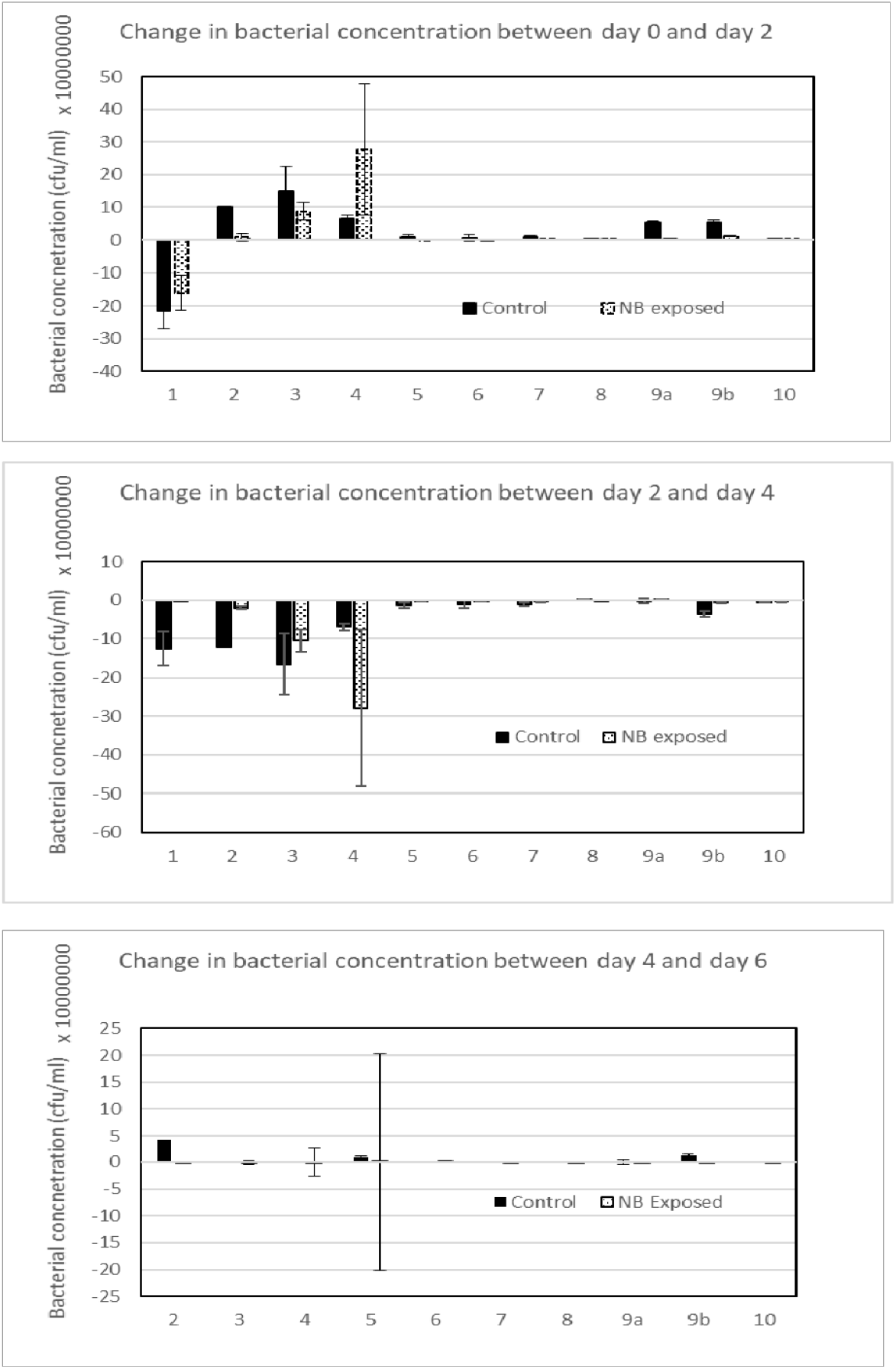
Relative difference in bacterial concentration between two day intervals for all experiments. A positive number indicates an increase in the bacterial concentration over the two-day period, a negative number indicates a decrease in the bacterial concentration over the two-day time period, and a number around zero indicates no change. The latter could have happened because there was no growth or decline in the bacterial concentration or there were no bacteria present in the tanks, which was the case at the end of many of these experiments.

Our nested linear regression model, which considered results of all experiments suggested overall there was no significant difference between the change in bacterial concentration from Day 0 to Day 2 between treated and untreated tanks (*p* = 0.596). Variance of the standardized residuals were considered equal when we examined them across treatments ignoring experimental groups. The normal distribution of the model residuals suggested this assumption for parametric general linear models was not met. Data could not be transformed to meet these assumptions. Given the robustness of the general linear model, and the fact that the differences in bacterial concentration were so small, we report our analyses despite the non-normal model residuals. The results of our Kruskal-Wallis tests on the individual experimental data also suggested the change in relative bacterial concentration between day 0 and day 2 for treated and untreated tanks was not significantly different for any experiments (Table 2).

**TABLE 2.**
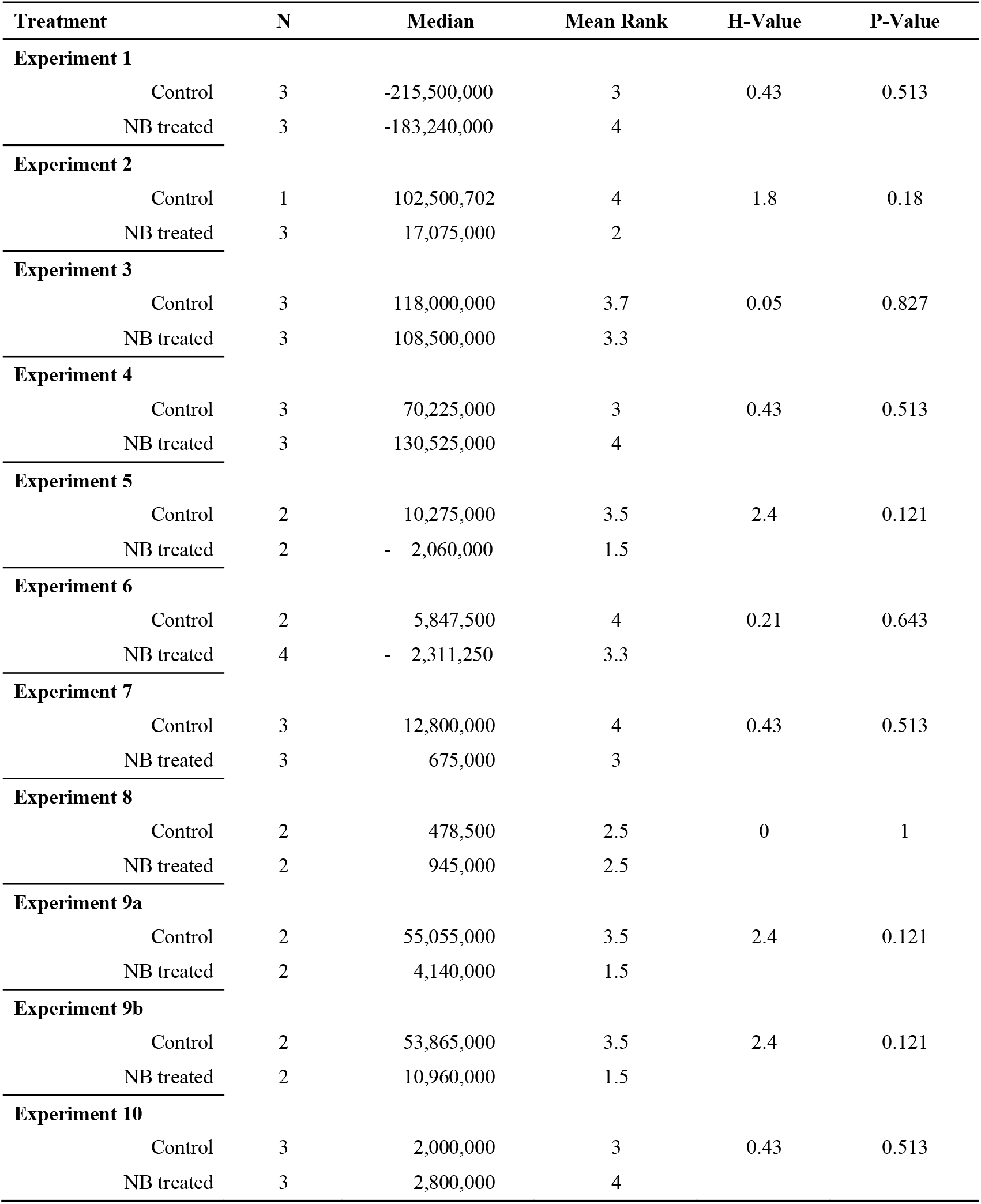
Results of Kruskal-Wallis analysis of the difference in bacterial concentration between day 0 and day 2 for individual experiments.

| Treatment | N | Median | Mean Rank | H-Value | P-Value |
| --- | --- | --- | --- | --- | --- |
| **Experiment 1** |  |  |  |  |  |
| Control | 3 | -215,500,000 | 3 | 0.43 | 0.513 |
| NB treated | 3 | -183,240,000 | 4 |  |  |
| **Experiment 2** |  |  |  |  |  |
| Control | 1 | 102,500,702 | 4 | 1.8 | 0.18 |
| NB treated | 3 | 17,075,000 | 2 |  |  |
| **Experiment 3** |  |  |  |  |  |
| Control | 3 | 118,000,000 | 3.7 | 0.05 | 0.827 |
| NB treated | 3 | 108,500,000 | 3.3 |  |  |
| **Experiment 4** |  |  |  |  |  |
| Control | 3 | 70,225,000 | 3 | 0.43 | 0.513 |
| NB treated | 3 | 130,525,000 | 4 |  |  |
| **Experiment 5** |  |  |  |  |  |
| Control | 2 | 10,275,000 | 3.5 | 2.4 | 0.121 |
| NB treated | 2 | - 2,060,000 | 1.5 |  |  |
| **Experiment 6** |  |  |  |  |  |
| Control | 2 | 5,847,500 | 4 | 0.21 | 0.643 |
| NB treated | 4 | - 2,311,250 | 3.3 |  |  |
| **Experiment 7** |  |  |  |  |  |
| Control | 3 | 12,800,000 | 4 | 0.43 | 0.513 |
| NB treated | 3 | 675,000 | 3 |  |  |
| **Experiment 8** |  |  |  |  |  |
| Control | 2 | 478,500 | 2.5 | 0 | 1 |
| NB treated | 2 | 945,000 | 2.5 |  |  |
| **Experiment 9a** |  |  |  |  |  |
| Control | 2 | 55,055,000 | 3.5 | 2.4 | 0.121 |
| NB treated | 2 | 4,140,000 | 1.5 |  |  |
| **Experiment 9b** |  |  |  |  |  |
| Control | 2 | 53,865,000 | 3.5 | 2.4 | 0.121 |
| NB treated | 2 | 10,960,000 | 1.5 |  |  |
| **Experiment 10** |  |  |  |  |  |
| Control | 3 | 2,000,000 | 3 | 0.43 | 0.513 |
| NB treated | 3 | 2,800,000 | 4 |  |  |

From Day 2 to Day 4 almost all experiments had a decline in bacterial levels in both their control and their NB exposed tanks, and by Day 6 all experiments with the exception of a few studies had very low concentrations of bacteria in both their control and the exposed tanks (Figure 2) there was also very little change in the concentration between these time points (Figure 3).

In addition, our flask experiment (# 11) with *S. agalactiae*, only detected differences in bacterial concentrations from 10 ^9.5^ cfu·ml-1 to 10 ^8^ cfu·ml-1 in all flasks (positive controls and single-dose nanobubble exposed) over the course of one week (exp. #11 in Figure 2). Water temperature was always slightly higher in tanks with NB than in control tanks. Once we introduced chillers or ice bottles during the nanobubble procedure we reduced this effect slightly (Table 1). Dissolved oxygen was also slightly higher when air nanobubbles were used in the tanks (Table 1).

## 4. DISCUSSION

Using several independent experiments, we evaluated air nanobubbles to determine the intrinsic disinfection properties of this technology. Three research teams, using similar laboratory protocols evaluated this technology had inconsistent results on bacterial replication. The summary of these experiments provides evidence that using **air** nanobubbles to disinfect water from aquatic pathogens in aquaculture would likely not be adequate to control disease. Despite the high concentration of air nanobubbles introduced daily to the relatively small volumes of water in experimental tanks, these bubbles had relatively little effect on the concentration of bacteria in the water. Although there were a few experiments where there appeared to be a decline in the concentration of bacteria in the nanobubble treated tanks, the 10 to 20-fold reduction in cfu·ml-1 (Figure 2) (i.e. a reduction in bacteria from 10 ^7^ to 10 ^6^ or 10 ^6^ to 10 ^5^ cfu·ml-1) was not statistically significant nor would it be biologically significant for reducing disease outbreaks in an aquaculture setting. We had expected a much greater reduction in bacterial concentration associated with nanobubbles given the promising literature on this technology (Gurunga et al., 2016).

In many experiments the concentrations of bacteria in the experiments usually increased during the first few days (Figures 2 and 3). This was most likely due to the enrichment media (TSB or NB), which was added to the tanks along with the bacteria, at the start of the experiment. However, in most of our experiments, bacteria naturally declined over time in both the control and the nanobubble exposure groups, unless additional media was added to the tanks, suggesting the reduction in bacteria had less to do with the nanobubbles and more to do with an environmental nutrient deficiency in both the control and the treated tanks. The use of green water for future studies with the addition of a higher level of nutrients may help alleviate this experimentally induced bacterial crash in 50 L tanks after 4 days. Despite the fact that we were not successful at maintaining a favourable environment for bacterial growth over a long period, we can conclude over a short period of time (i.e. 2 days), where we did not have issues with bacterial survival, we could not consistently and significantly reduce the relative concentration of bacteria using high concentrations of air nanobubbles. At most, and only in a few experiments we may have reduce bacteria by 10 or 20-fold, but this could not be replicated in many of the experiments.

We predominantly assessed Gram-negative bacteria in fresh water in these experiments (i.e. *Aeromonas* spp.). We believed, based on the fact that nanobubbles have a larger zeta-potential (charge) in lower salinity water (Li et al., 2013; Hu and Xia, 2018), and Gram-negative bacteria have a thinner peptidoglycan layer than Gram-positive bacteria (Mai-Prochnow et al., 2016). Gram-negative bacteria in fresh water may be more prone to damage from free OH· radicals generated from nanobubbles. Interestingly the only experiments that showed any potential minor effect of nanobubbles were those using fresh water and *Aeromonas* spp (Figure 2). Regardless, despite an excessive concentration of air nanobubbles the impact in freshwater with *Aeromonas* spp. was minimal. Our findings on the bactericidal effects of air nanobubbles on several types of bacteria suggest that this technology, was not sufficient to reduce bacteria to a level that would prevent or mitigate bacterial disease outbreaks.

The concentration of bacteria required to trigger disease in aquaculture settings depends on the pathogenicity of the bacterial species, but in general in laboratory studies researchers use a target concentration of 10 ^4^ or greater to induce disease (O’Toole et al., 1996; Evans et al., 2000; Harikrishnan et al., 2003; AlYahya et al., 2018). We were not able to achieve a reduction in bacterial concentration below 10 ^5^ by day 2 in any of our experiments even when we started with a concentration of bacteria at 10 ^6^ cfu·ml-1. Although there were a few experiments that appeared to have a slight reduction in bacteria over the first few days, this could not be consistently repeated across the research institutions with relatively similar laboratory set ups. Even with the addition of the chillers, which reduced the increase in temperature of NB treated tanks that could have counteracted the bactericidal effects of nanobubbles, we still had bacterial growth in tanks exposed to nanobubbles in two out of the 4 experiments (Experiment # 8 & 9a,b). It is possible with more replicates the relative small decrease in bacterial concentration observed in some of the experiments would have been statistically significant, but the effect would still not be sufficient to be biologically significant. Perhaps substituting air for a gas such as ozone or oxygen would improve the disinfection properties of nanobubbles as has been suggested by several researchers (Kurita et al., 2017; Imaizumi et al., 2018, Jhunkeaw et al., 2020).

It is difficult to ensure that nanobubble generators produce the appropriate size of bubbles to elicit the unique physical properties of nanobubbles, because it is difficult to detect these bubbles without specialized equipment. In this study, we evaluated the bubbles that our nanobubble generator produced before the experiments were initiated using a NanoSight LM10. Because this machine is not able to distinguish bubbles from solid particles, we had to test the nanobubble size using distilled ultraclean water. Under these conditions bubble size was consistent across samples, and as other studies have indicated the nanobubbles appeared to persist over time (Hu and Xia, 2018; Wang et al., 2019). Results from our nanoparticle analyser studies suggest that the nanobubble generator produced approximately 10 ^11^ bubbles per minute and ∼ 70 % of these bubbles were nanobubble size (< 150 nm). Given these figures, we roughly estimate that when we operated our generator for 30 min in 100 L of water, we had at least 2.0 × 10 ^7^ nanobubbles per ml of water in the tanks. In addition, we introduced nanobubbles every day to the tanks.

Because nanobubbles are relatively stable in water (Figure 1), the concentration of nanobubbles in the tanks would have increased over time. Although the concentration of nanobubbles likely varied somewhat at the different research sites, given the amount of time we operated the nanobubbler and the size of the tanks, it is unlikely that low concentrations of nanobubbles was the reason that we did not observed significant reductions in bacterial concentration in our experiments. We used a high concentration of nanobubbles in the tanks to ensure we were not missing any potential effect of air nanobubbles on bacteria. We did this because the flask experiment (designated as Experiment 11), which was conducted using water treated with a single dose of nanobubbles did not produce a change in the bacterial concentration (Figure 2 Experiment #11). It is important to note that the concentration of nanobubbles in these experiments would be difficult to achieve in a commercial aquaculture setting.

## 5. CONCLUSION

In this series of independent experiments, we explored the disinfection potential of air nanobubbles on pathogenic bacteria in water. We determined that the application of these nanobubbles, did not significantly or consistently alter the level of bacteria in the water compared to the control tanks in our experiments. This technology may disinfect water more effectively when ozone is used to create the nanobubbles, as has been reported by other researchers (Kurita et al., 2017; Imaizumi et al., 2018, Jhunkeaw et al., 2020). This is being explored further.

## ACKNOWLEDGEMENTS

This work was carried out with financial support from UK government – Department of Health and Social Care (DHSC), Global AMR Innovation fund (GAMRIF) and the International Development Research Center (IDRC), Ottawa, Canada.

## DISCLAIMERS

The views expressed herein do not necessarily represent those of IDRC or its Board of Governors. None of the authors have any conflict of interests.

## AUTHOR CONTRIBUTIONS

All authors were actively involved in the described study. The laboratory in Shenzhen China had four researchers : Qianjun Huang (student), Hong Liu (Lab coordinator), Jose Domingos (post doctoral research) and Sophie St-Hilaire (Project leader). They were responsible for conducting experiment # 1-7. The laboratory in Vietnam had four researchers: Phan Thi Van (research lead), Nguyen Huu Nghia (student), Pham Thai Giang (technician), Pham The Viet (technician). This laboratory conducted experiment # 8,9,11. The laboratory in Thailand had two researchers Ha Thanh Dong (head of the laboratory) and Nareerat Khongcharoen (student). This laboratory conducted experiment # 10. The project manager (Jose Domingos) assisted in writing and analyzing the metadata with the help of Prof. Sophie St-Hilaire.

## Notes

### Competing Interest Statement

The authors have declared no competing interest.

